# Brain Organoid Computing for Artificial Intelligence

**DOI:** 10.1101/2023.02.28.530502

**Authors:** Hongwei Cai, Zheng Ao, Chunhui Tian, Zhuhao Wu, Hongcheng Liu, Jason Tchieu, Mingxia Gu, Ken Mackie, Feng Guo

**Affiliations:** Department of Intelligent Systems Engineering, Indiana University Bloomington, IN, 47405, United States; Department of Industrial and Systems Engineering, University of Florida, Florida, 32611, Gainesville, United States; Center for Stem Cell and Organoid Medicine (CuSTOM), Division of Pulmonary Biology, Division of Developmental Biology, Cincinnati Children’s Hospital Medical Center, OH, 45229, Cincinnati, United States; University of Cincinnati School of Medicine, OH, 45229, Cincinnati, United States; Gill Center for Biomolecular Science, Department of Psychological and Brain Sciences, Indiana University Bloomington, IN, 47405, United States

**Keywords:** Artificial Intelligence (AI), Brain-inspired Hardware, Neuromorphic Computing, Biological Neural Networks, Brain Organoids

## Abstract

Brain-inspired hardware emulates the structure and working principles of a biological brain and may address the hardware bottleneck for fast-growing artificial intelligence (AI). Current brain-inspired silicon chips are promising but still limit their power to fully mimic brain function for AI computing. Here, we develop *Brainoware*, living AI hardware that harnesses the computation power of 3D biological neural networks in a brain organoid. Brain-like 3D *in vitro* cultures compute by receiving and sending information via a multielectrode array. Applying spatiotemporal electrical stimulation, this approach not only exhibits nonlinear dynamics and fading memory properties but also learns from training data. Further experiments demonstrate real-world applications in solving non-linear equations. This approach may provide new insights into AI hardware.

## Introduction

Artificial intelligence (AI) is reshaping the future of human life across various real-world fields such as industry, medicine, society, and education^1^. The remarkable success of AI has been largely driven by the rise of artificial neural networks (ANNs), which process vast numbers of real-world datasets (big data) using silicon computing chips ^2, 3^. However, current AI hardware keeps AI from reaching its full potential since training ANNs on current computing hardware produces massive heat and is heavily time-consuming and energy-consuming ^4–6^, significantly limiting the scale, speed, and efficiency of ANNs. Moreover, current AI hardware is approaching its theoretical limit and dramatically decreasing its development no longer following ‘Moore’s law’^7, 8^, and facing challenges stemming from the physical separation of data from data-processing units known as the ‘von Neumann bottleneck’^9, 10^. Thus, AI needs a hardware revolution^8, 11^.

A breakthrough in AI hardware may be inspired by the structure and function of a human brain, which has a remarkably efficient ability, known as natural intelligence (NI), to process and learn from spatiotemporal information. For example, a human brain forms a 3D living complex biological network of about 200 billion cells linked to one another via hundreds of trillions of nanometer-sized synapses^12, 13^. Their high efficiency renders a human brain to be ideal hardware for AI. Indeed, a typical human brain expands a power of about 20 watts, while current AI hardware consumes about 8 million watts to drive a comparative ANN^6^. Moreover, the human brain could effectively process and learn information from noisy data with minimal training cost by neuronal plasticity and neurogenesis,^14, 15^ avoiding the huge energy consumption in doing the same job by current high precision computing approaches^12, 13^. The human brain fuses data storage and processes within biological neural networks (BNNs),^16–18^ naturally avoiding any ‘von Neumann bottleneck’ issues. Learning from BNNs, pioneering attempts have been made to develop high-efficiency and low-cost neuromorphic chips (e.g., memristor)^11, 19, 20^ that store previously experienced current or/and voltage in internal states and enable short-term memory ^21–23^. So far, neuromorphic chips have been used for various applications in computer vision,^24–26^ speech recognition,^27, 28^ and others ^29, 30^. However, current neuromorphic chips can only mimic partial features of brain functions, and there are tremendous needs to improve their processing capability for real-life uncertainty and energy efficiency.

Brain organoids are *in vitro* 3D aggregates derived through the self-organization and differentiation of human pluripotent stem cells that become brain-like tissues with a remarkable ability to recapitulate aspects of a developing brain’s structure and function^31–34^. Herein, we develop *Brainoware*, living AI hardware that harnesses the computation and on-chip learning of 3D biological neural networks in a brain organoid (**Fig. 1a**). *Brainoware* processes and learns from spatiotemporal information through the neuroplasticity of the brain organoid. Compared to current 2D in vitro neuronal cultures and neuromorphic chips, *Brainoware* may provide new insights for AI computing, because brain organoids can provide 3D BNNs with remarkable advances in complexity, connectivity, neuroplasticity, and neurogenesis, all accomplished with low energy consumption and fast learning.

**Fig. 1.**
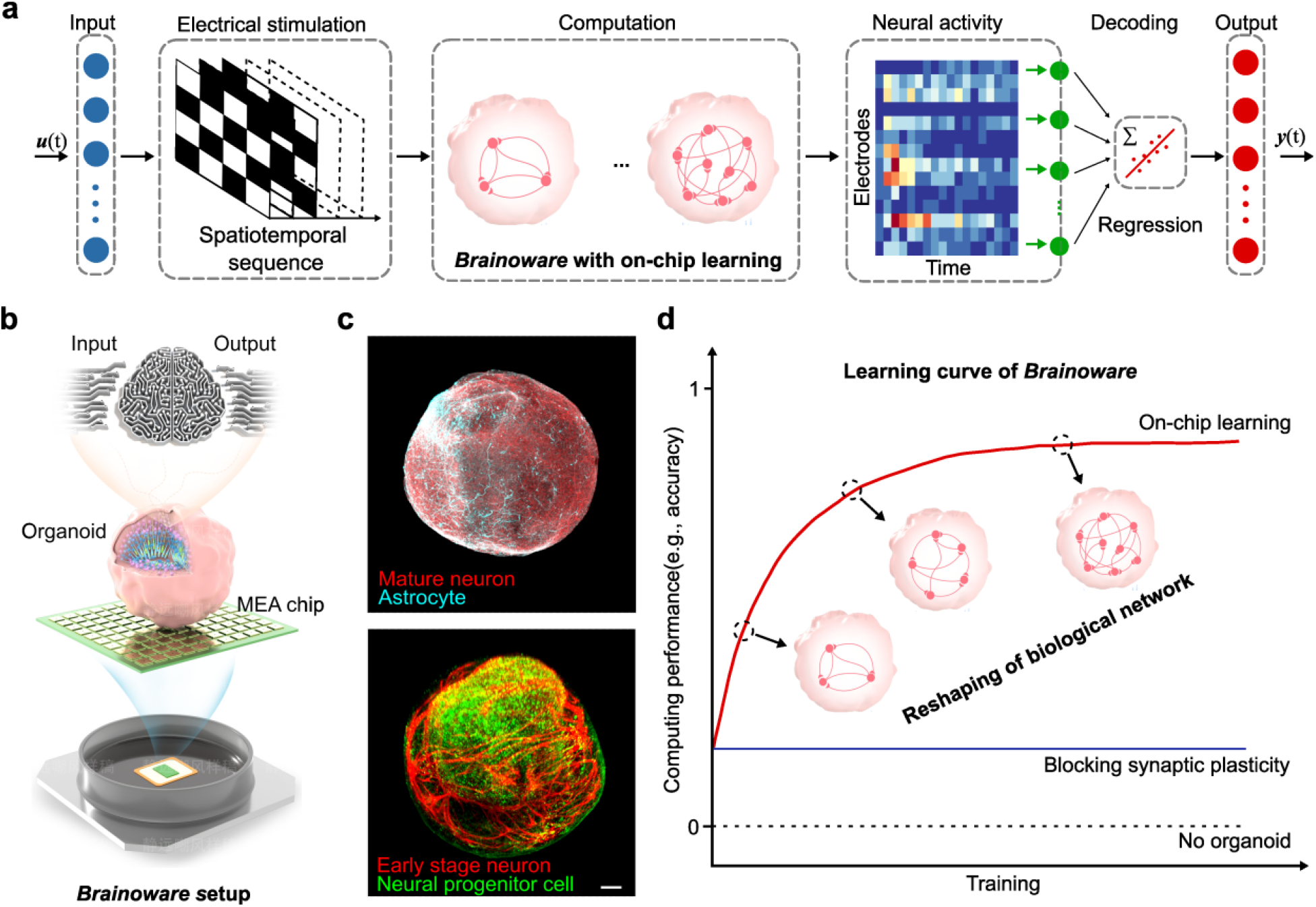
*Brainoware* with on-chip learning for AI computing. **a,** Schematics of reservoir computation with on-chip learning using *Brainoware*. **b,** A schematic paradigm of *Brainoware* setup that mounts a brain organoid onto a multielectrode array for receiving inputs and sending outputs. **c**, Whole-mount immunostaining of cortical organoids showing complex 3D neuronal networks with various brain cell identities (e.g., mature neuron, MAP2; astrocyte GFAP; neurons of early differentiation stage, TuJ1; and neural progenitor cells, SOX2). **d**, Schematic diagram demonstrating on-chip learning of *Brainoware* via reshaping of the biological neural network during training, and the inhibition of on-chip learning after synaptic plasticity is blocked. (Scale bar = 100 μm)

## Results

### *Brainoware* with on-chip learning for AI computing

We constructed *Brainoware* by mounting a functional brain organoid onto a multielectrode array (MEA) (**Fig. 1b**). The mature human brain organoid for *Brainoware* was characterized by various brain cell identities (e.g., early-stage and mature neurons, astrocytes, neuron progenitor cells), and early development brain-like structures (e.g., including ventricular zones and subventricular zones) for the formation, function, and maintenance of complex 3D neuronal networks (**Fig. 1c**), as well as network electrical activity. The mature organoid neural networks (ONNs) could receive inputs via external electrical stimulation and send outputs via evoked neural activity, offering a functional basis for AI computing. As a proof-of-concept application, we implemented *Brainoware* as an AI hardware to solve real-world problems. In a conventional AI computing hardware, input signals can be mapped into higher dimensional computational spaces through the dynamics of a hardware. Given input signals, the output of hardware is used as features for a simple “readout function” to perform a computational task, e.g., time-series analysis. While the conventional computing chips are fixed, the readout function is trained to map the feature values generated by the hardware to the desired labels of the data. Different from conventional AI computing hardware with fixed physical reservoirs, *Brainoware* uses mature human brain organoids as dynamic, living network to conduct “unsupervised learning” to adapt to the signals and perform “feature engineering”. Via these living reservoirs, the time-dependent inputs can be converted to spatiotemporal sequences of electric stimulation and then projected into computational spaces as ONNs. The output signals, as neural activities, can be effectively utilized via the pre-trained readout function for various tasks (**Fig. 1a**). Moreover, through training using spatiotemporal sequences of electrical stimulation, *Brainoware* can improve its computing performance and demonstrate on-chip learning ability by synaptic plasticity. This is possible because *Brainoware* responds to the electric stimulations with changes in neuronal connections, and neuroplasticity of the organoids, enabling dynamic reshaping of ONNs. If synaptic plasticity is blocked, computing performance is maintained by *Brainoware* (but on-chip learning halts) (**Fig. 1d, Fig. 3d**). In the next experiments, *Brainoware* was demonstrated to exhibit unique properties of a physical reservoir and successfully conduct some real-world tasks with limited training data at low energy and computing cost.

**Fig. 2.**
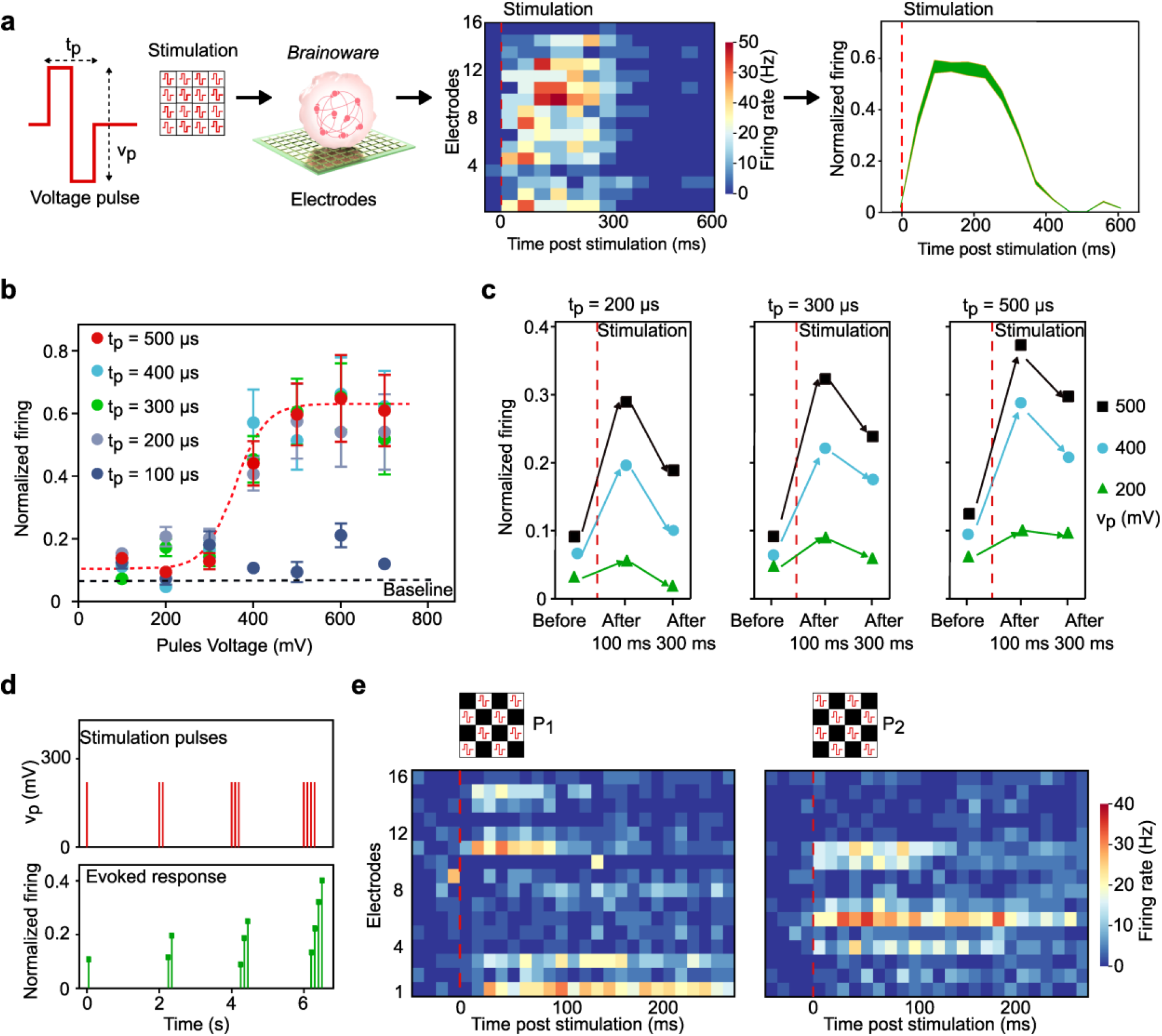
Characterization of hardware property. **a,**Evoked response (raster plot and post-stimulation histogram) upon single bipolar voltage pulse stimulation. **b,** Evoked normalized firing upon pulses with different pulse times (t_p_) and pulse voltage (v_p_) (mean ± s.e.m., n =6, independent experiments). The red fitting curve (a sigmoid function) indicates the nonlinear activity, while the black dashed line marks the spontaneous activity. **c,** Evoked normalized firing before and after, 100 ms or 300 ms, from the end of single pulse stimulation (mean ± s.e.m., n = 4, independent experiments). **d,** Representative memristor-liked responses to a stream of pulses (v_p_ = 200mV, t_p_ = 300 μs). **e,** Distinct raster plots evoked by two complementary spatial patterns (P1 and P2) of stimulation pulses (vp = 500 mV, tp = 500 μs).

**Fig. 3.**
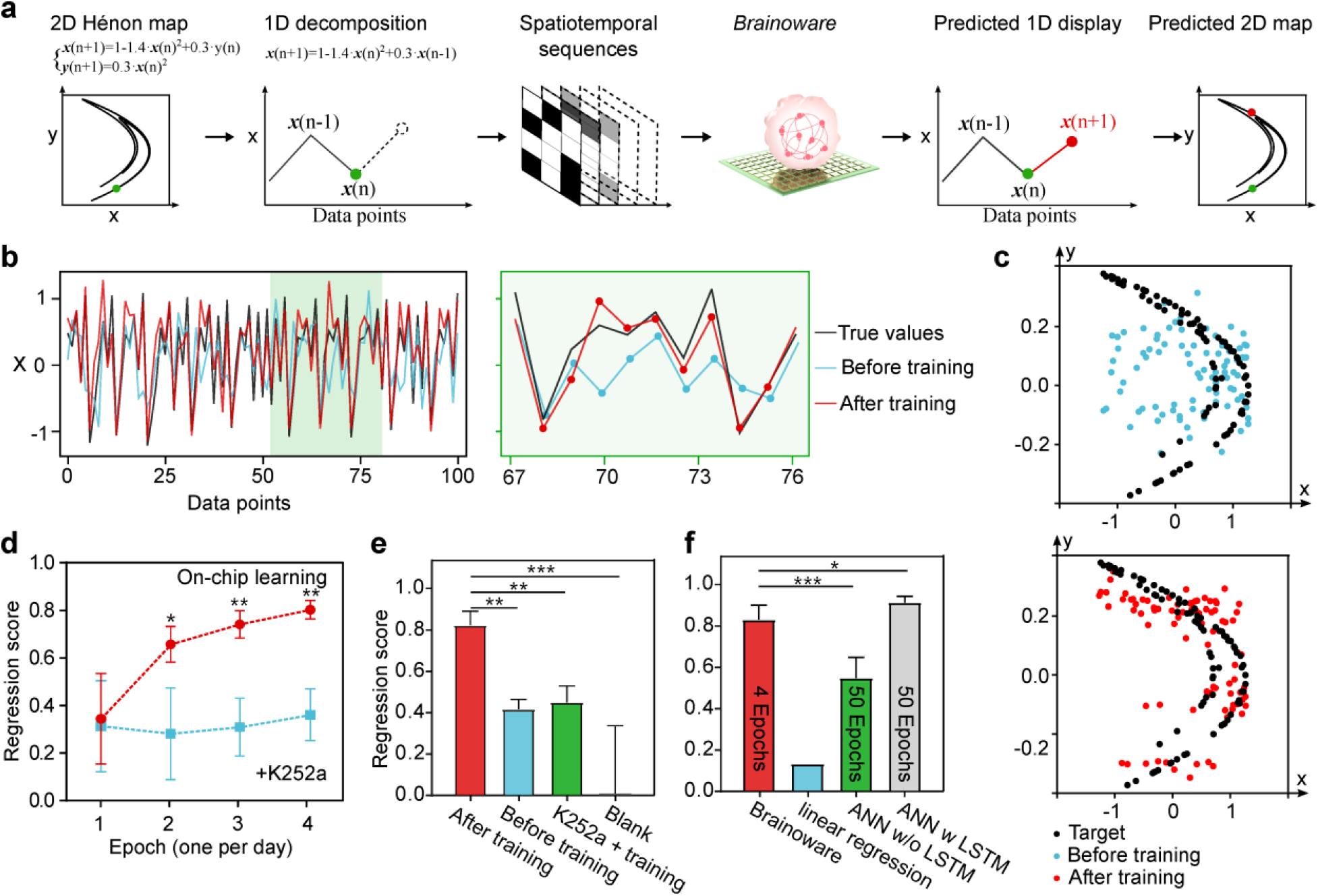
Solving a non-linear chaotic equation. **a,** Workflow of predicting an Hénon map. **b,** Predicted *X* values using *Brainoware* before (blue) and after training (red) vs ground true *X* value. **c,** Predicted 2D maps using *Brainoware* before (blue) and after (red) training vs ground true 2D map. **d,** Learning curves of *Brainoware* over training epochs, where the red or blue curves show *Brainoware* with naïve organoids or organoids treated with a CaMKII blocker K252a (to block synaptic plasticity) before training (mean ±s.e.m., n =4, independent experiments). **e,** Learning activity of *Brainoware* under different conditions with the naïve organoids (before training), the organoids after training (after training), the organoids treated with K252a during training (K252a + training), and without organoids (Blank) (mean ± s.e.m., n = 5 for each group, independent experiments). **f,** Performance comparison of *Brainoware* with linear regression, artificial neural network (ANN) with or without long short-term memory (LSTM) unit. The number denotes the training epochs of each group (mean ± s.e.m., n = 5 for each group, independent experiments).

### Characterization of hardware property

Before applying *Brainoware* to AI computing tasks, we characterized and demonstrated its basic implementation as a AI hardware. We tested the physical reservoir properties of *Brainoware* such as nonlinear dynamics, fading memory (or short-term memory), and spatial information processing by checking the response of ONNs to bipolar voltage pulse stimulations with different pulse time (t_p_) and voltage (v_p_). For example, while applying electrical stimulation pulses to *Brainoware*, the evoked neuronal activity (raster plot) was recorded, and the post-stimulation histogram was calculated and plotted (**Fig. 2b**). We demonstrated that *Brainoware* exhibited a representative nonlinear response to the pulse voltage (**Fig. 2b**). After applying a single voltage pulse stimulation (t_p_ >= 200 μs), the evoked mean normalized firing rate of *Brainoware* (over 200 ms post-stimulation) to the pulse voltage can be fitted with a sigmoid function, in accordance with the nonlinear activation function of ANNs. While applying a single voltage pulse stimulation with short pulse times (t_p_ < 200 μs), the evoked normalized firing of *Brainoware* with the same organoid was only around the baseline of its spontaneous activity. Next, we tested the fading memory of *Brainoware* by applying pulses with different pulse times and voltages. The evoked normalized firing of *Brainoware* before and after 100 ms or 300 ms from the end of single pulse stimulation was obtained (**Fig. 2c**). Pulses with longer duration and higher voltage were responsible for stronger evoked response and slower relaxation dynamics. Importantly, the nonlinear response and fading dynamics of ONNs can be well controlled by precisely adjusting the stimulation parameters. Moreover, we also demonstrated the combination of these two properties within *Brainoware*. After the application of four individual trains of pulses (v_p_ = 200mV, t_p_ = 300 μs), *Brainoware* showed both the accumulation and decay dynamic responses (**Fig. 2d**). Multiple pulses at short intervals (50 ms) within a pulse train were responsible for the gradual increase of evoked responses and the delay of relaxation dynamics, in accordance with the dynamic response of a memristor, a typical reservoir computing hardware. Furthermore, we demonstrated the capability of *Brainoware* to process spatial information. The spatial information was converted into spatial patterns of simulation pulses (v_p_ = 500mV, t_p_ = 500 μs) such as two 4 x 4 spatial patterns (P1 and P2). The distinct raster plots of *Brainoware* with the same organoid were evoked by these two complementary patterns and showed the active storage and gradual loss of different spatial information over time (**Fig. 2e**), indicating spatial information processing rather than stimulation artifacts.

### Solving a non-linear chaotic equation

We further applied *Brainoware* to predict a Hénon map, which is a typical non-linear dynamic system with chaotic behavior. This time-series task was implemented into *Brainoware* using a brief workflow as shown in **Fig.3a**. A 2D Hénon map was first converted into a 1D decomposition, converted into the spatiotemporal sequences of bipolar voltage pulses, and then sent to the MEA electrodes for stimulating *Brainoware*. Using a read-out algorithm for decoding the ONN activity, *Brainoware* harnessed its reservoir computation and on-chip learning to learn from the input spatiotemporal pulses and predict the Hénon map. Experiments were conducted to predict the Hénon map (X_n+1_ value) by feeding *Brainoware* with spatiotemporal pulses encoded with X_n_ value. 1D decomposition (**Fig.3b**) and 2D displacement (**Fig.3c**) of predicted Hénon maps were experimentally obtained from *Brainoware* with the same organoid before and after four training epochs (one epoch per day, and each epoch encoded with a Hénon map dataset of 200 data points). Compared with the theoretical output (ground truth, black), the after-training condition (red) showed better-predicted results than the before-training condition (blue). Next, the learning curves of *Brainoware* to solve this non-linear chaotic equation were measured over epochs (**Fig.3d**). The regression score of *Brainoware* in predicting X_n+1_ value was used to evaluate its learning ability to solve the Hénon map. Interestingly, *Brainoware* increased the regression score (detailed calculation in SI) from 0.356 ±0.071 (with the naïve organoids on Day 1) to 0.812 ±0.043 (the same organoids after four training epochs). While treated with a calcium/calmodulin-dependent protein kinase II (CaMKII) blocker, K252a, to block activity-dependent synaptic plasticity,^35^ the negative control group only slightly improved their regression score from 0.31 ±0.072 to 0.385 ±0.063 over the same training procedures. The results indicated that the learning activity of *Brainoware* was dependent on neural plasticity. Furthermore, experiments were performed to measure the on-chip learning activities of *Brainoware* under different conditions (**Fig. 3e**). Only an MEA chip and culture medium (blank) was tested to have a regression score of 0, emphasizing that *Brainoware* cannot compute without the living organoids. We further compared *Brainoware* with representative machine learning algorithms such as ANN on solving the Hénon map of the same data size (**Fig. 3f**). Linear regression (decoding algorithm of *Brainoware*) could barely solve this problem, showing a regression score close to 0. *Brainoware* significantly outperformed ANN without a long short-term memory (LSTM) unit. *Brainoware* (with 4 training epochs) showed slightly lower accuracy than ANN with LSTM (each with 50 training epochs), decreasing training times by >90%.

## Discussion

We have demonstrated a new class of living hardware for AI computing that harnesses the computational power of living neural networks of human brain organoids. Human brain organoids have unique characteristics and the ability to self-organize and form functional BNNs for developing brain-inspired AI hardware. Moreover, ONNs may have the necessary complexity and diversity to mimic a human brain, inspiring more sophisticated and human-like AI systems. Furthermore, due to the high plasticity and adaptability of organoids, *Brainoware* has the flexibility to change and reorganize in response to electrical stimulation, highlighting its ability to learn and adapt over training, necessary for developing AI systems. As living brain-like AI hardware, this approach may naturally address the time-consuming, energy-consuming, and heat production challenges of current AI hardware. We further demonstrated that this approach exhibits physical reservoir properties such as nonlinear dynamics, fading memory, and spatial information processing. We also implemented it in real-world applications such as solving non-linear equations. We found that this approach learns from training data by reshaping the neuronal connections of ONNs.

There are several limitations and challenges for the current *Brainoware* approach. One technical challenge is the generation and maintenance of organoids. Despite the successful establishment of various protocols, current organoids are still suffering from high heterogeneity, low generation throughput, and various viabilities. Moreover, it is critical to properly maintain and support organoids for harvesting their computational power. Recent engineering efforts on optimizing organoid differentiation and growth conditions and manipulating their microenvironments may provide approaches for the high-throughput generation and maintenance of standardized organoids^36^. *Brainoware* uses flat and rigid MEA electrodes for interfacing with organoids, which are only able to stimulate/record a small number of neurons on the organoid surface. There is a tremendous need to develop methods such as brain-machine interfaces, soft electrodes, and other neural implants for interfacing the whole organoid with AI hardware and software ^37, 38^, allowing for the exchange of information and the manipulation of their activity from a greater number of neurons. Another technical challenge is the management and analysis of data. The encoding and decoding of temporospatial information to and from *Brainoware* still need optimization by improving data interpretation, extraction, and processing from multiple sources and modalities. Moreover, large amounts of data may be generated by this new AI hardware, requiring the development of new algorithms and methods for analyzing and visualizing the data.

In sum, we developed a prototype of brain-inspired living AI hardware with on-chip learning ability. We also demonstrated their real-world applications to speech recognition and solving non-linear equations. This approach may provide new insights into AI hardware.

## Methods

### Generation and characterization of organoids

Cortical organoids were generated from human pluripotent stem cells following a protocol that we adapted from the reported protocols^39^. All the handling and culture of stem cells and organoids followed the guidelines of the WiCell Institute and Indiana University Biosafety Committee. Detailed protocols for the development and characterization of organoids are included in the supplementary materials.

## Acknowledgment

This work was funded with support from the National Institute of Health Awards (DP2AI160242, R01DK133864., and U01DA056242). We also acknowledge Indiana University Imaging Center (NIH1S10OD024988-01).

## Author contributions

F.G. and H.C. conceived the study and designed experiments. H.C., Z.A., C.T., and Z.W. performed the experiment. H.C., H.L., J.T., M.G., and K.M. analyzed the data. All authors read and provided feedback on the manuscript.

## Competing interests

Authors declare that they have no competing interests.

## Notes

### Competing Interest Statement

The authors have declared no competing interest.

